# Beyond the Hype: Deep Neural Networks Outperform Established Methods Using A ChEMBL Bioactivity Benchmark Set

**DOI:** 10.1101/168914

**Authors:** Eelke B. Lenselink, Niels ten Dijke, Brandon Bongers, George Papadatos, Herman W.T. van Vlijmen, Wojtek Kowalczyk, Adriaan P. IJzerman, Gerard J.P. van Westen

**Author notes:** Authors contributed equally to this work (Eelke B. Lenselink, Gerard J.P. van Westen). Present address: GlaxoSmithKline, Medicines Research Centre, Gunnels Wood Road, Stevenage, Herts SG1 2NY, UK. Author email addresses: EB Lenselink, N ten Dijke, B Bongers, G Papadatos, HWT van Vlijmen, W Kowalczyk, AP IJzerman, GJP van Westen.

## Abstract

The increase of publicly available bioactivity data in recent years has fueled and catalyzed research in chemogenomics, data mining, and modeling approaches. As a direct result, over the past few years a multitude of different methods have been reported and evaluated, such as target fishing, nearest neighbor similarity-based methods, and Quantitative Structure Activity Relationship (QSAR)-based protocols. However, such studies are typically conducted on different datasets, using different validation strategies, and different metrics.

In this study, different methods were compared using one single standardized dataset obtained from ChEMBL, which is made available to the public, using standardized metrics (BEDROC and Matthews Correlation Coefficient). Specifically, the performance of Naive Bayes, Random Forests, Support Vector Machines, Logistic Regression, and Deep Neural Networks was assessed using QSAR and proteochemometric (PCM) methods. All methods were validated using both a random split validation and a temporal validation, with the latter being a more realistic benchmark of expected prospective execution.

Deep Neural Networks are the top performing classifiers, highlighting the added value of Deep Neural Networks over other more conventional methods. Moreover, the best method (‘DNN_PCM’) performed significantly better at almost one standard deviation higher than the mean performance. Furthermore, Multi task and PCM implementations were shown to improve performance over single task Deep Neural Networks. Conversely, target prediction performed almost two standard deviations under the mean performance. Random Forests, Support Vector Machines, and Logistic Regression performed around mean performance. Finally, using an ensemble of DNNs, alongside additional tuning, enhanced the relative performance by another 27% (compared with unoptimized DNN_PCM).

Here, a standardized set to test and evaluate different machine learning algorithms in the context of multitask learning is offered by providing the data and the protocols.

## 1 Introduction

The amount of chemical and biological data in the public domain has grown exponentially over the last decades [1-3]. With the advent of ChEMBL, computational drug discovery in an academic setting has undergone a revolution [4, 5]. Indeed, the amount of data available in ChEMBL is also growing rapidly (**Supplementary figure 1**). Yet data availability and data quality still pose limitations [6]. Public data is sparse (on average a single compound is tested on two proteins) and prone to experimental error (on average 0.5 log units for IC_50_ data) [6, 7]. To make full use of the potential of this sparse data and to study ligand-protein interactions on a proteome wide scale, computational methods are indispensable as they can be used to predict bioactivity values of compound-target combinations that have not been tested experimentally [8-10].

In order to compare our work to established target prediction methods, we framed the original problem as a classification task by labeling compound-protein interactions as ‘active’ or ‘inactive’. Data explored here contains pChEMBL values, which represent comparable measures of concentrations to reach half-maximal response / effect / potency / affinity transformed to a negative logarithmic scale. The threshold at which molecules are labeled ‘active’ determines the fraction of data points belonging to the ‘active’ compound class. If this is set at 10 μM (pChEMBL = 5) as is done frequently in literature, almost 90% of the extracted ChEMBL data is an ‘active’ compound making it the default state (**Supplementary figure 2**) [10, 11].

Hence, predictions out of the model will likely be ‘active’. Such a high fraction of active compounds is not in accordance with what is observed experimentally. Moreover, in an experimental context, model output should ideally lead to identification of compounds with affinity higher than 10 μM to make most efficient use of costly experimental validation. Based on these considerations, we chose to set the decision boundary at 6.5 log units (approximately 300 nM), defining interactions with a log affinity value larger than or equal to 6.5 as ‘active’ compounds. At this boundary, the distribution between ‘active’ and ‘inactive’ compounds is roughly 50% (**Supplementary figure 2**). For reference, a model using the 10 μM threshold and a Naïve Bayesian classifier was included in this study which could be seen as a baseline.

Furthermore, as was touched upon above, public data can have relatively large measurement errors, mostly caused by the data being generated in separate laboratories by different scientists at different points in time with different assay protocols. To make sure that bioactivity models are as reliable as possible, we chose to limit ourselves to the highest data quality available in ChEMBL (**Supplementary figure 3**) using only confidence class 9 data points were used. A common misconception in literature is that the confidence class as implemented in ChEMBL is interpreted as quality *quantification* rather than *classification* (i.e., the higher the confidence, the better, using data points confidence class higher than 7). Yet, this is not always true as the confidence represents different classes (i.e., ‘homologous protein assigned’ as target versus ‘direct target assigned’). Hence some confidence classes are not compatible with each other for the goal pursued by a method. An example of a confidence class 8 assay is: CHEMBL884791 and an example of a class 9 assay is CHEMBL1037717. Both compound series have been tested on the Adenosine A_2A_ receptor but in the former case it was obtained from bovine striatal membrane and the latter explicitly mentions human Adenosine A_2A_ receptors. In the current work, we chose consistently class 9 (see the recent paper by the ChEMBL team on data curation and methods for further details) [6].

It has been shown that compounds tend to bind to more than one target, moreover compounds have sometimes been tested active on multiple proteins [12, 13]. This activity spectrum can be modeled using (ensembles of) binary class estimators, for instance by combining multiple binary class RF models (**Figure 1**). Another strategy is to assemble one model with all targets, which can be done in various ways. With multiclass QSAR (MC), it can be predicted if a compound is active based on the probability of belonging to the active target class versus the inactive target class for a given target; each compound-target combination is assigned ‘active’ or ‘inactive’ (**Figure 1**).

**Figure 1:**
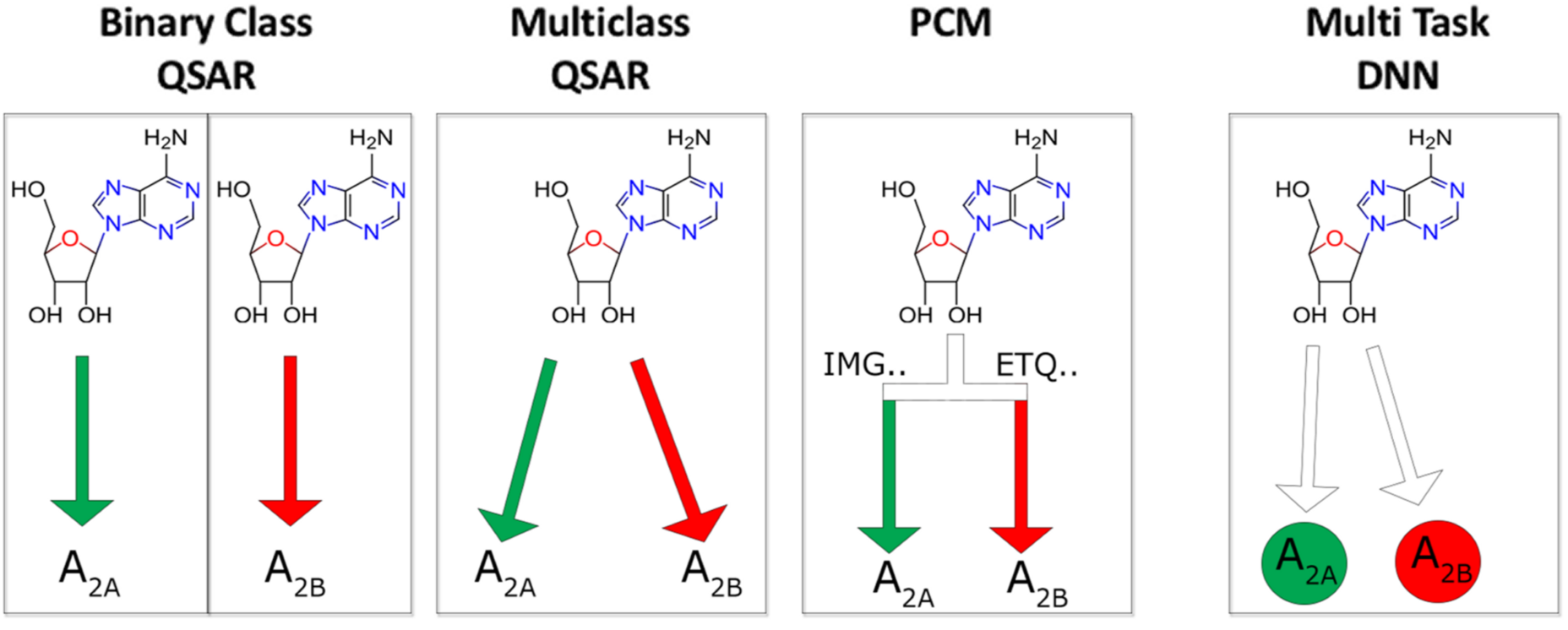
Differences between methods for modeling bioactivity data exemplified by the ligand adenosine which is more active (designated as ‘active’) on the adenosine A2A receptor, than on the A2B receptor (‘inactive’, using PChEMBL > 6.5 as a cutoff). With binary class QSAR, individual models are constructed for every target. With multiclass QSAR one model is constructed based on the different target labels (A2A_active, A2B_inactive). With PCM one model is constructed where the differences between proteins are considered in the descriptors (i.e. based on the amino acid sequence). With multiclass DNN a single output node is explicitly assigned to each target.

Yet another approach is to apply machine learning algorithms with added protein descriptors, commonly known as proteochemometrics (PCM) [14, 15]. Targets are represented in the data in the form of target descriptors and this is combined with compound descriptors. Hence, instead of determining the activity of a compound, the activity of a compound / protein pair (**Figure 1**) is determined. Explicitly quantifying this protein similarity allows models to make predictions for targets with no or very little bioactivity data but for which a sequence is known. Moreover, explicit protein features allow interpretation of both the important protein and ligand features from the validated model. The relationships between structure and biological activity in these large pooled datasets are non-linear and best modeled using non-linear methods such as RF or Support Vector Machines (SVM) with non-linear kernels. Alternatively, when linear methods are used cross-terms are required that account for the non-linearity [16].

Several of our models benchmarked here are multitask, however for simplicity we grouped the different methods on underlying machine learning algorithm. Nevertheless, multitask learning has been shown to outperform other methods in bioactivity modeling and the reader is referred to Yuan *et al*. for a more in depth analysis [17].

Another non-linear method is Deep Neural Networks (DNNs), which have recently gained traction being successfully applied to a variety of artificial intelligence tasks such as image recognition, autonomously driving cars, and the GO-playing program AlphaGO [18, 19]. Given their relative novelty they will be introduced here in relation to our research but the reader is referred to LeCun *et al*. for a more extensive introduction of the subject.[19] Deep Neural Networks have many layers allowing them to extract high level features from the raw data. DNNs come in multiple shapes but here we focus only on fully connected networks, i.e., networks where each node is connected to all the nodes in the preceding layer.

In feed forward networks (such as implemented in the current work) information moves from the input layer to the output layer through ‘hidden’ layers (which can be one layer to many layers). Each hidden node applies a (usually) non-linear response function to a weighted linear combination of values computed by the nodes from the preceding layer. By this, the representation of the data is slightly modified at each layer, creating high level representations of the data. The behavior of the network is fully determined by the weights of all connections. These weights are tuned during the training process by an optimization algorithm called backpropagation to allow the network to model the input-output relation. The major advantage of DNNs is that they can discover some structure in the training data and consequently incrementally modify the data representation, resulting in a superior accuracy of trained networks. In our research, we experimented with several scenarios, such as training as many networks as the number of targets or just one network with as many output nodes as the number of targets. (**Figure 1**).

DNNs have been applied to model bioactivity data previously; in 2012 Merck launched a challenge to build QSAR models for 15 different tasks [20]. The winning solution contained an ensemble of single-task DNNs, several multi-task DNNs, and Gaussian process regression models. The multi-task neural networks modeled all 15 tasks simultaneously, which were subsequently discussed in the corresponding paper [20]. Later (multi-task) DNNs have also been applied on a larger scale to 200 different targets [21], tested in virtual screening [22], and was one of the winning algorithms of the Tox21 competition [23]. Recently different flavors of neural networks also have shown to outperform random forests on various, diverse cheminformatics tasks [24]. Hence, DNNs have demonstrated great potential in bioactivity modeling, however they have not been tested in a PCM approach to the best of our knowledge. Therefore, they have been included in our research as this technique may become the algorithm of choice for both PCM and QSAR.

Summarizing, we perform a systematic study on a high quality ChEMBL dataset, using two metrics for validation Mathews Correlation Coefficient (MCC) and Boltzmann-Enhanced Discrimination of ROC (BEDROC). The MCC was calculated to represent the global model quality, and has been shown to be a good metric for unbalanced datasets [25]. In addition, BEDROC represents a score that is more representative of compound prioritization, since it is biased towards early enrichment [26]. The BEDROC score used here (α=20) corresponds to 80% of the score coming from the top 8%.

We compare QSAR and PCM methods, multiple algorithms (including DNNs), the differences between binary class and multi-label models, and usage of temporal validation (effectively validating true prospective use). We used both open- and closed-source software, we provide the dataset and predictions, PP protocols to generate the dataset, and scripts for the DNNs in the Supplementary Information and on the internet (see section 4.10 and a DOI after acceptance).

Hence, the current work contributes to the literature by providing not only a standardized dataset available to the public, but also a realistic estimate of the performance that published methods can currently achieve in preclinical drug discovery using public data.

## 2 Results and discussion

### 2.1 Random split partition

Firstly, the accuracy of all algorithms was estimated with help of the random split method (**Figure 2**). Models were trained on 70% of the random split set and then validated on the remaining 30%. Validation of the classifier predictions on a multi-target dataset such as this one can be done either based on all predictions in a single confusion matrix or by calculating a confusion matrix per target and subsequently using the mean value. Both methods provide relevant information, thus we followed both and show the mean and the standard error of the mean (SEM) obtained from these two sets of experiments. Multiclass Random Forests were trained but omitted due to their poor performance (as can be seen in **Supplementary table 1** and **2**). For LR or SVM no PCM models were completed. An LR PCM model would require cross terms due to linearity, which would make a direct comparison impossible. Training of SVM PCM models was stopped after running for over 300 hours. Since results for Python (with scikit-learn) and Pipeline Pilot (PP, using R-statistics) were comparable in most cases, the results reported here are for the Python work with the PP results in the Supplementary Information. The exception is the 10 μM NB model, trained in PP which is our baseline model. Individual results for all methods are reported in the Supplementary information. (**Supplementary table 2)**.

**Figure 2:**
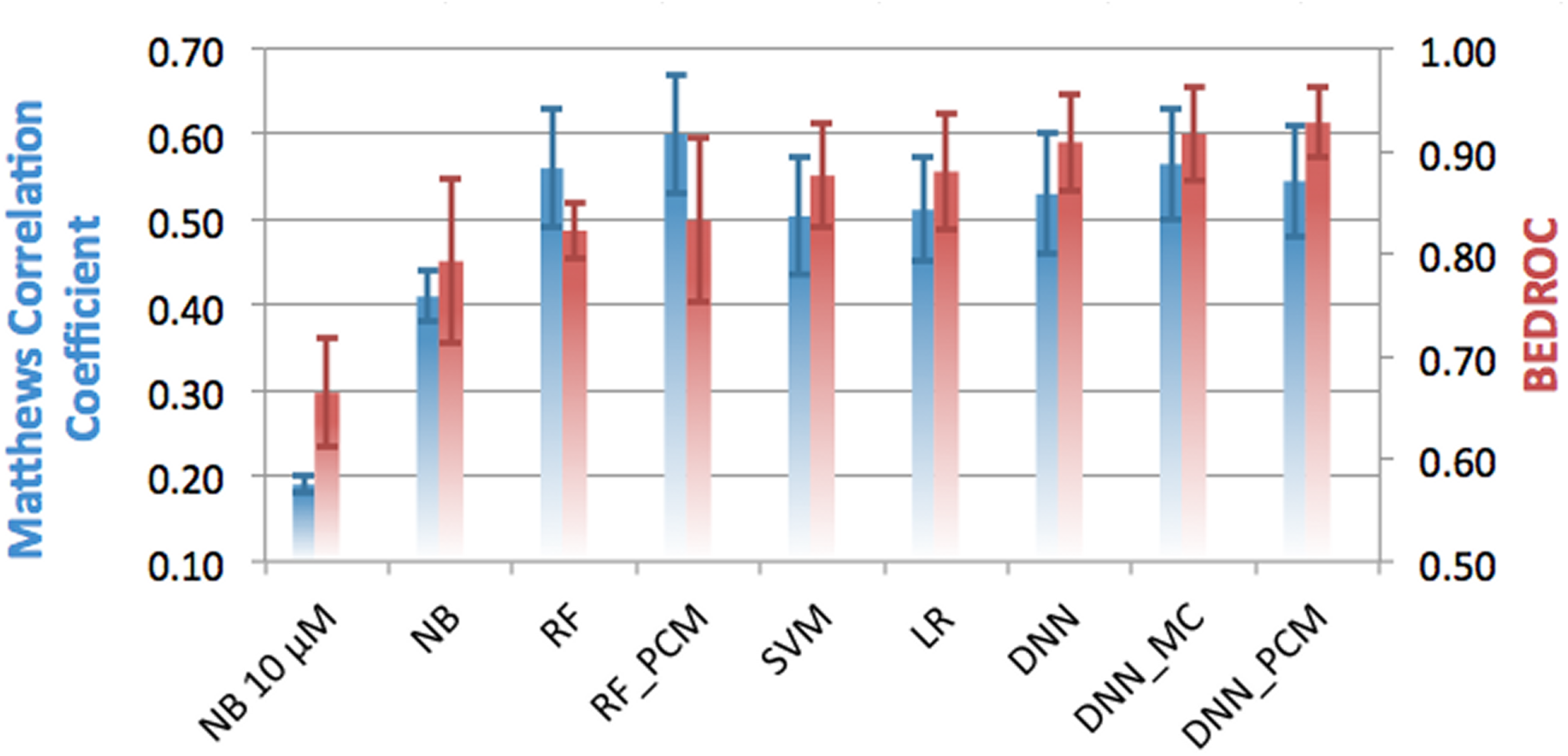
Performance of the different methods in the random split validation, grouped by underlying algorithm and colored by metric used. On the left y-axis, and in blue the Matthews Correlation Coefficient is shown, while on the right y-axis and in red the BEDROC (α = 20) score is shown. Default, single class algorithms are shown, and for several algorithms the performance of PCM and multi-class implementations is shown. Error bars indicate SEM. Mean MCC is 0.49 ( ±0.04) and mean BEDROC is 0.85 (± 0.03).

The average MCC of all algorithms is 0.49 (± 0.04), underlining the predictive power of most methods. The mean BEDROC was 0.85 (± 0.03), which corresponds with a high early enrichment. The performance of all DNNs are well above the average performance, both in terms of MCC and BEDROC. The best method overall is the DNN_MC with an MCC of 0.57 (± 0.07), and a BEDROC score of 0.92 (± 0.05). DNN_PCM is performing slightly worse (MCC of 0.55 ± 0.07), but slightly better in terms of BEDROC (0.93 ± 0.03) and the DNN follows (MCC of 0.53 (± 0.07) and BEDROC of 0.91 (± 0.05)).

The worst performing method is the NB 10 μM (MCC of 0.19 ± 0.01 and BEDROC 0.66 ± 0.05), where NB (using the 6.5 log units activity threshold) performs around the mean of all performing methods (MCC of 0.41 ± 0.03 and BEDROC 0.79 ± 0.08). Indeed, using an activity threshold of 6.5 log units appears to improve performance. Surprisingly logistic regression performed above the average (MCC of 0.51 ± 0.06 and BEDROC 0.88 ± 0.06). However, large differences were observed between logistic regression in Pipeline Pilot and Python, most likely due to the fact that the latter uses regularization (**Supplementary table 1**).

Overall it is found that high/low MCC scores, typically also corresponded with high/low BEDROC scores with some exceptions. Most notably was the RF_PCM model which was the best performing model in terms of MCC (MCC of 0.60 ± 0.07), but underperformed in terms of BEDROC (BEDROC 0.83 ± 0.08). Moreover, judged on MCC the QSAR implementation of RF outperforms SVM (MCC of 0.56 ± 0.07 versus 0.50 ± 0.07). Yet, based on the BEDROC, SVM outperforms the RF model (BEDROC 0.88 ±0.05 versus 0.82 ± 0.03). Based on this we pose that SVMs are better in predicting top ranking predictions, but RFs are better in predicting negative predictions.

While these results look encouraging, it should be noted that in a random splitting scenario all data points (measured activities of protein-compound combinations) are considered separate entities. Hence, members of a congeneric compound series from a given publication can be part of the test set while the remaining are part of the training set (see methods - validation partitioning). Therefore, this method is expected to give an optimistic estimate of model performance; for a more representative performance estimate, a more challenging validation is exemplified below.

### 2.2 Temporal split partition

In the temporal split, training data was grouped by publication year rather than random partitioning (**Figure 3**). All data points originating from publications that appeared prior to 2013 were used in the training set, while newer data points went into the validation set. Using temporal split we aim to minimize the effect that members of a congeneric chemical series are divided over training and test set. Temporal split has previously been shown to be a better reflection of prospective performance, than other validation schemes [27].

**Figure 3:**
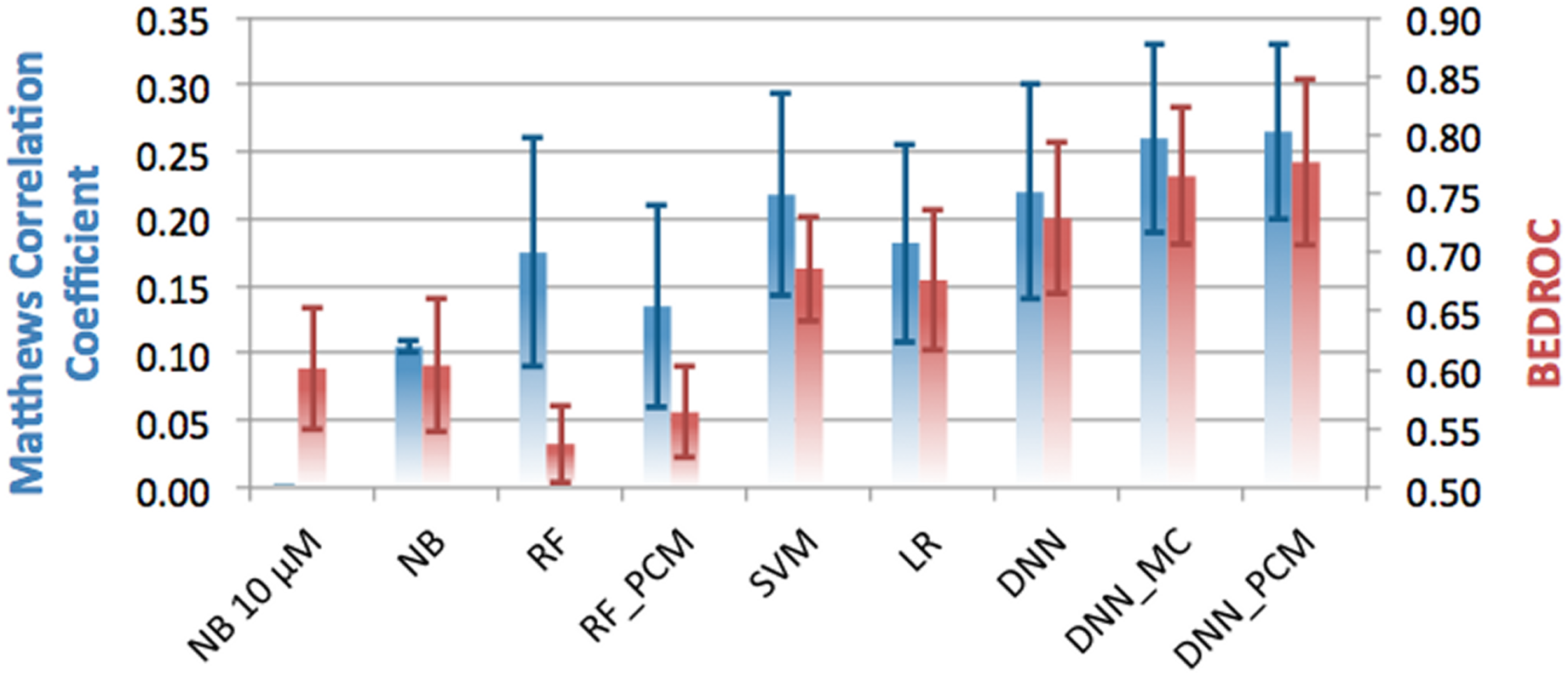
Performance of the different methods in the temporal split validation, grouped by underlying algorithm and colored by metric used. On the left y-axis, and in blue the Matthews Correlation Coefficient is shown, while on the right y-axis and in red the BEDROC (α = 20) score is shown. Default, single class algorithms are shown, and for several algorithms the performance of PCM and multi-class implementations is shown. Error bars indicate SEM. MeanMCC is 0.17 (±0.03) and mean BEDROC is 0.66 (± 0.03)

All methods performed worse than on the random split benchmark. The average MCC dropped to 0.18 (± 0.03) from 0.49 (± 0.04) in the random split with a similar picture for the BEDROC 0.66 (± (0.03) from 0.85 (± 0.03). A paired t-test p-value < 0.01 for both MCC and BEDROC was obtained, confirmed that this is indeed a significantly more challenging form of validation (**Supplementary table 2**).

Large differences between methods are observed, for instance the RF model in terms of MCC is performing around the average, but both RF and RF_PCM underperform in terms of early enrichment (BEDROC 0.54 ± 0.03 and 0.56 ± 0.04 versus the mean 0.66 ± 0.03). SVM (MCC of 0.22 ± 0.07 and BEDROC 0.69 ± 0.07) performed in between the DNNs and RF models. Both NB 10 μM and NB underperform based on MCC and BEDROC. Finally, all DNNs outperformed the other methods both in terms of MCC (0.22 ± 0.08 - 0.27 ± 0.07) and even more so in terms of BEDROC (0.73 ± 0.06 – 0.78 ± 0.07). For the DNN_PCM, we found that for targets with few data points in the training set, the PCM models were able to extrapolate predictions (**Supplementary figure 4**).

Summarizing, the lower performance observed here is more in line with the performance that can be expected from a true prospective application of these types of models. It has been suggested in literature that also temporal splitting is not ideal, but it still provides a more challenging form of validation and better than leaving out chemical clusters [27]. Hence, this make temporal split validation a better way to validate computational models. Yet, in addition to raw performance, training time is also of importance.

### 2.3 Run time

Quick training models allow for easy retraining when new data becomes available, models that require a long time are not readily updated, making their maintenance a tradeoff. It was found that on our hardware most models could be retrained in under 10 hours. This training time corresponds with an overnight job (**Supplementary table 3**).

One point should be visited, NB in Pipeline Pilot was considerable slower than the NB trained in scikit-learn (20 minutes in scikit-learn compared with 31 hours, **Supplementary table 3**). This is caused by the calculation of the background scores (see methods for details) as was done previously [28]. Calculation of z-scores requires the prediction of all ligand – protein interactions in the matrix and is a lengthy procedure regardless of the high speed of NB. As can be seen, the NB 10 μM models do not suffer this penalty (as they do not use z-score calculation) and are hence the fastest.

When we compare training time with MCC and BEDROC, we observe that training times do not directly correlate with the quality of the model. In both cases a weak trend is observed between performance and training time (R^2^ 0.25 and 0.38 respectively, **Supplementary figure 5**). It should be noted that RF can be trained in parallel (on CPUs) leading to a speedup in wall clock time but that there is a saturation around 40 cores [29]. In addition, the parallel implementation requires installation of additional third party packages such as ‘foreach’ for the R implementation [30]. In scikit-learn this works more efficiently, however, in both cases running in parallel increases memory consumption. Note that a GPU version of RF (CUDAtrees) was published in 2013 but this package is no longer maintained (abandoned November 2014). Hence, while RF can be optimized, this is not as straightforward as in DNN. Still, it should be noted that the GPU-implementation of the DNN speeds up the calculation about ~150 times when compared with the CPU-implementation (benchmarked on a single core); this makes GPUs a definite requirement for the training of DNNs.

### 2.4 Ranking the various methods

To accurately compare the methods, 4 z-scores were calculated for each method and metric within the experiments (random split MCC, random split BEDROC, temporal split MCC, and temporal split BEDROC, **Table 1** and **Figure 4**). Herein DNNs are found to be the best algorithm, and have the most consistent performance. For instance, for DNN the best model (DNN_PCM) has an average z-score of 0.96 (± 0.19), compared to -0.69 (± 0.04) for the best NB model and a slightly better -0.21 (± 0.41) for the best RF model. Moreover, the three best methods based on the average z-score are all DNNs, which are subsequently followed by SVM (0.32 ± 0.09). Furthermore, the DNNs perform the best in all types of validations in terms of BEDROC and MCC, with a single exception (the random split MCC where RF_PCM is the best as can be observed in bold in **Table 1**). To confirm whether the observed differences were actually statistically significant, the following tests were performed: the paired Student’s T-Test (determine whether the means of two groups differ), the F-Test (determine whether the variances of groups differ), Wilcoxon Rank test (determine whether sets have the same median), and the Kolmogorov-Smirnov test (determine whether two samples come from the same random distribution). Results can be found in the supporting information **Supplementary tables 4 - 7**, here the results will be summarized.

**Table 1.** Overview of the performance of the benchmarked methods. Z-scores are shown for all methods for both types of splitting andfor both MCC and BEDROC. In bold the best performance for a given machine learning algorithm per column is highlighted. See main text for further details.

| Method | MCC<br>Random | BEDROC<br>Random | MCC<br>Temporal | BEDROC<br>Temporal | Average | SEM |
| --- | --- | --- | --- | --- | --- | --- |
| NB 10 uM | -2.41 | -2.22 | -2.07 | -0.67 | -1.84 | 0.40 |
| NB | -0.65 | -0.66 | -0.81 | -0.64 | -0.69 | 0.04 |
| RF | 0.56 | -0.30 | 0.02 | -1.41 | -0.28 | **0.41** |
| RF_PCM | **0.88** | -0.17 | -0.46 | -1.10 | -0.21 | **0.41** |
| SVM | 0.11 | 0.36 | 0.53 | 0.30 | 0.32 | 0.09 |
| LR | 0.17 | 0.40 | 0.11 | 0.19 | 0.22 | 0.06 |
| DNN | 0.32 | 0.75 | 0.56 | 0.79 | 0.60 | 0.11 |
| DNN_MC | 0.60 | 0.85 | 1.03 | 1.20 | 0.92 | 0.13 |
| DNN_PCM | 0.44 | **0.98** | **1.09** | **1.33** | **0.96** | 0.19 |

**Figure 4:**
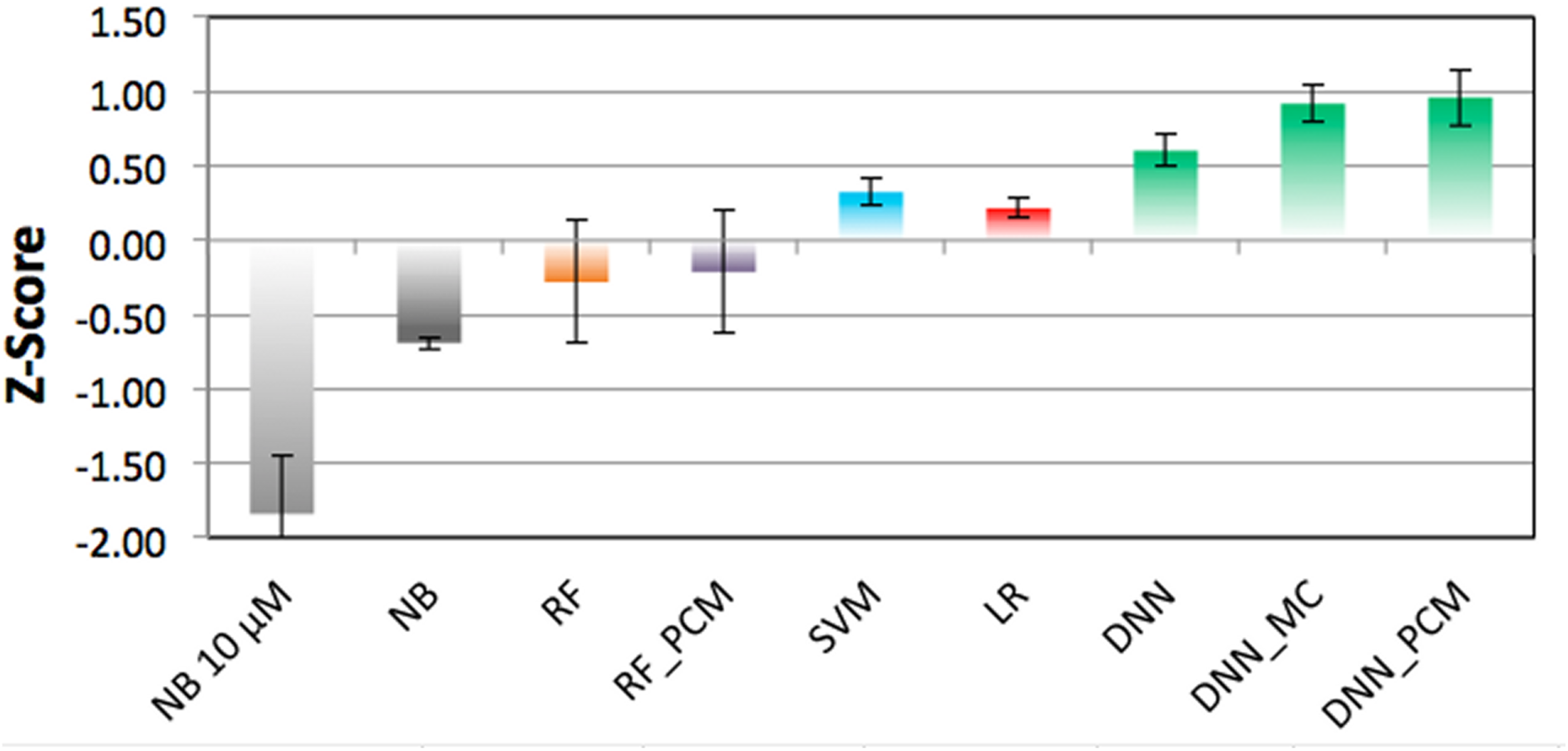
Comparison of the mean z-scores obtained by the different methods. Bars are colored by method and error bars indicate SEM, best performance is by the DNN (0.96 ±0.19, 0.92 ±0.13, and 0.60 ±0.11 respectively), followed by SVM (0.32 ±0.09), LR (0.22 ±0.06), RF (-0.21 ±0.41 and-0.28 ±0.41), and finally NB (-0.69 ±0.04 and-1.84 ±0.40).

For the Student’s T-Test DNN_PCM and DNN_MC are shown to significantly differ from all other methods with a p-value < 0.05 except for RF and RF_PCM, where the p-values are 0.05, 0.06 (2 times), and 0.07. DNN is seen to differ significantly from NB 10 μM, NB, and LR. Likewise, NB 10 μM differs significantly from all other methods with the p-value < 0.05 with the exception of NB, where the p-value is 0.06. It can hence be concluded that there are little differences in performance using RF, SVM, and LR, whereas the use of NB is significantly worse than the rest and usage of DNN leads to significantly better results.

In the variances (F-Test) less significant differences are found. NB 10 μM differs significantly from NB, SVM, LR. Similarly, NB differs significantly from RF, RF_PCM, and DNN_PCM. RF and RF_PCM differ significantly from SVM, LR, DNN (with the exception of the pair RF_PCM - DNN which has a p-value of 0.06). Hence in general variance in SVM and LR differs significantly from NB and RF, whereas between the other methods not really significant differences exist.

The results of the Wilcoxon and Kolgomorov-Smirnov test were very similar to each other. For both, the differences between SVM, LR, DNN, DNN_MC, DNN_PCM on one hand and both NB 10 μM and NB on the other hand are significant. Secondly, in both DNN_MC differs significantly with RF, SVM, and LR. Finally, DNN_PCM differs significantly with LR in both. In general it can be concluded that NB and RF methods differ significantly from other methods and DNN_MC differs from most (based on the methods median value and whether samples come from the same random distribution).

In conclusion, here it was shown that DNN methods generally outperform other algorithms and that this is a statistically significant result. However, we used DNN as is and it should be noted that there is room for improvement by (among other things) inclusion of more information and training of hyper parameters which will be further explored in the next section.

### 2.5 Exploring the potential of DNNs

An additional reason that DNNs were of interest in this study is the fact that they can process more information without a high penalty in training time. Because in general DNNs are quite sensitive to the choice of hyper parameters, we explored a number of different parameters through a grid search based exploration of model parameter space. For this we varied the architecture of the networks, ranging from one layer of 1000 hidden nodes (very shallow), to two layers of 2000 and 1000 nodes (shallow), to the default settings (4000, 2000, 1000 nodes) and the deepest network used here (8000, 4000, 2000 nodes).

In addition to the number of nodes we varied the dropout which represents the percentage of nodes that are dropped randomly during the training phase, a technique to prevent overfitting [31]. By default (as used above), there is no dropout in the input layer and 25% on the hidden layers. However, in the increased dropout scenario 25% dropout is introduced in the input layer and in the higher layers increased to 50%.

Thirdly, the usage of more extensive compound descriptors was investigated (up to 4096 bits and additional physicochemical descriptors) which was not possible with the RF and NB models due to computational restraints.

Finally, differences between PCM, MC, and QSAR were investigated. To test all these different combinations mentioned above the maximum number of epochs was decreased from 2000 to 500 (see methods - machine learning methods - neural networks). These settings were validated on the temporal split because it represents a more realistic and more challenging scenario as shown in section 2.3.

The predictive performance of all the DNNs are summarized in **Figure 5**, while performance of individual models is shown in **Supplementary figure 6**. Improvements are expressed in the percentage of increase over the baseline performance as demonstrated by the DNN_PCM covered in section 2.2 (temporal split). Based on these results the following can be concluded. First of all, performance using a longer bit string is better (9% improvement for 4096 bits with extra features compared to the baseline), with on the low end of the performance spectrum the 256 bits that were used prior to this optimization (average decrease of 2% in the grid search). This intuitively makes sense as the shorter fingerprints contain less information. Moreover, it could be that distinct chemical features computed by the fingerprint algorithm hash to the same bit. In that case a specific bit could represent the presence of multiple features, which is more likely to happen with a shorter fingerprint. Furthermore, out of the models trained with 256 bits descriptors for ligands, the PCM DNN consistently outperformed the others, likely due to the fact that PCM profits from the added protein features also containing information.

**Figure 5:**
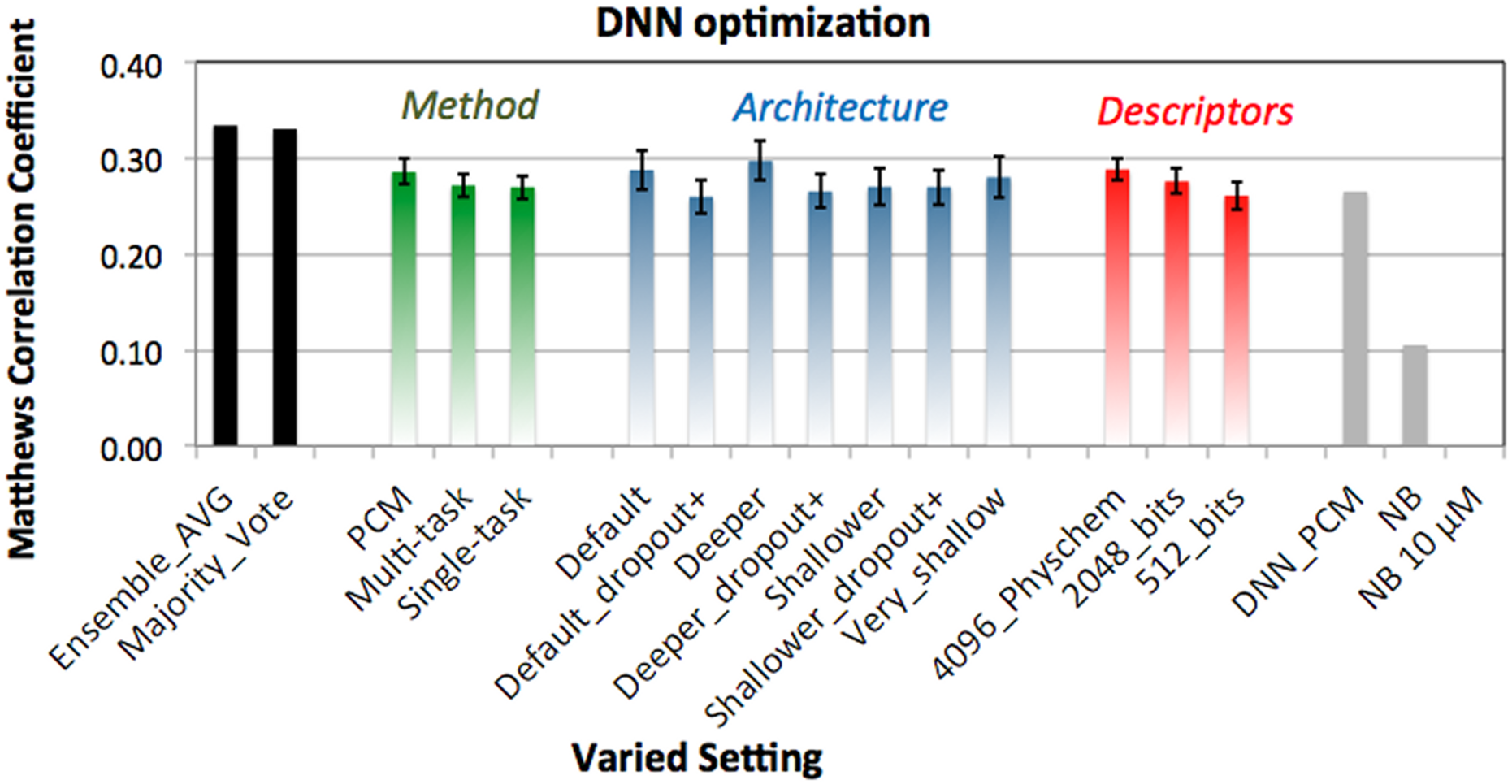
Average performance of the individual DNN grouped per method, architecture and descriptors. Average value is shown for all models trained sharing a setting indicated on the x-axis, error bars represent the SEM of that average. Black bars on the left represent the ensemble methods (average value and majority vote). Grey bars on the right indicate the previous best performing DNN (DNNPCM), NB with activity cut-off at 6.5 log units and z-score calculation, and default NB with activity cut-off at 10 μM. We observed PCM to be the best way to model the data (green bars), architecture 3 to be the best performing (blue bars), and usage of 4096 bit descriptors with additional physicochemical property descriptors to perform the best (red bars). Using ensemble methods further improves performance (black bars).

Of the three different DNNs, PCM slightly outperforms the other methods (average improvement 8%), although examples of both single and multi-task models are also found in the top performing methods (average increase 2% and 2% respectively). With regard to the architecture, deep and wide networks seem to perform best (e.g., architecture 3 with an average increase of 12%), although some of the shallow, multiclass and binary class networks (architecture 7) are also found in the top performing methods.

Overall it seems that increasing dropout leads to a poorer performance. Since dropout is a technique to prevent overfitting, a DNN can be considered as underfitted if dropout is too strict. This is confirmed by these results, as higher dropout rates and dropout on the visible layer (the fingerprint/feature layer) results in a drop of accuracy (1 versus 2 and 3 versus 4). Moreover, if all increased dropout results and normal dropout results are aggregated, increased dropout performs near identical to the baseline (0%) and the normal dropout architectures (on average) perform 7% better than baseline. Therefore, an option to be considered is a less aggressive dropout. Alternatively lowering the dropout percentage adaptively (during the training), similar to the learning rate would be an option too.

Finally, the best performance is observed by using an ensemble of the predictions from all models, for instance by using a majority prediction or an average vote (improvement 25% and 26%, black bars **Figure 5**). This improvement suggests that there is still room for further improvement and only the surface was scratched in this work. Indeed, ensembles of different machine learning methods, including neural networks have been used to achieve competitive results on bioactivity prediction tasks. More context will be discussed below.

### 2.6 Putting this work into context

As touched upon, ChEMBL has fueled a diverse array of publications and this discussion is limited to the most relevant and recent ones in the context of this paper. For instance Mervin *et al*. constructed a (Bernoulli) Bayesian model on both ChEMBL and PubChem data [32]. To balance the class sizes, a sphere exclusion algorithm was used to extract putative inactive data. A different threshold was used (10 μM, pChEMBL = 5) compared to the current study, and PubChem data in addition to the ChEMBL data was used. Also in the current work, it was found that inclusion of inactive molecules enhanced the performance for the Naive Bayes models (**Supplementary table 8**).

A later study by Lusci *et al*. that was performed on ChEMBL data release 13 benchmarked the performance of a number of different algorithms [33]. Similar to the current work the authors performed a temporal validation (on ChEMBL release 13) as a more realistic estimate of model performance. They also found that their method, potency-sensitive influence relevance voter (PS-IRV) outperformed other methods such as RF and SVM. However, here it is proposed that limiting the training set to high quality data with only the highest confidence from ChEMBL, leads to better performance. This is also corroborated by the AUC values obtained by Lusci *et al*. on their full set and the higher values obtained in the current work. IRV has been benchmarked before [34], and can be seen as an extension of K-nearest neighbors in a shallow neural network. In that study, random molecules were added (presumably inactive), in addition to the experimentally inactive molecules to boost results. Inclusion of more data, and more specifically inactive molecules is a line of future investigation we also aim to pursue.

Regarding the DNNs, the influence of network architecture has been studied before [20], where it was noted that the number of neurons especially impacts the performance of deeper structured networks. This corresponds to our observations where the deepest and widest network performed best. Further fine-tuning of the architecture might be worthwhile; in multi-task networks trained for the Tox21 challenge up to 4 layers with 16,384 units were used [23]. Additionally, it was found that multiclass networks outperformed binary class networks, and similar gains in performance were observed on the joint (multi-task) DNN published by Ma *et al*. [20]. This is also in line with our own results where DNN is seen to slightly improve over state of the art methods such as RF and SVM, but DNN_MC and DNN_PCM are demonstrated to really improve performance.

Finally, work by Unterthiner *et al*. demonstrated similar DNN performance [22]. Though the authors did not calculate the BEDROC, they obtained an AUC of 0.83 versus the here obtained AUC of 0.89 (**Supplementary table 1**). Interestingly, they found a worse NB performance (AUC 0.76 versus 0.81) compared to the current work [22]. This divergence is potentially caused by the fact that their dataset included lower quality ChEMBL data, which was the main reason for assembling the current benchmark dataset. Moreover, Unterthiner *et al*. used much larger feature input vectors, requiring ample compute power to use the non-DNN based algorithms. We have shown that we can achieve similar performance on a smaller dataset with fewer fingerprint features, suggesting that there is much room for improvement by hyperparameter optimization. Furthermore, Unterthiner *et al*. used a cost function weighted, based on the dataset size for every target. In our hands, experimentation choosing different weights inversely proportional to the target dataset size did not improve the performance of the models, however this can be further be explored. Finally, we have shown that usage of (simple) ensemble methods outperformed a single DNN alone, hence more sophisticated ensemble methods and inclusion of different models is a worthy follow up.

DNNs have also been applied with promising results to the prediction of Drug-Induced Liver Injury [35], although a different descriptor was used than the conventional fingerprints, i.e. directed acyclic graph recursive neural networks [36]. Similarly, convolutional networks were recently applied to molecular graphs, outperforming extended connectivity fingerprints (ECFP) [37]. Interestingly, contrary to ECFP, such graphs are directly interpretable [38]. This work was further extended to a diverse palate of different cheminformatics datasets, in MoleculeNet, where a lot of different architectures have been tested and compared [24]. Moreover, these methods are also publicly available in the form of the package DeepChem, a package for DNNs that is actively maintained. Future work will focus on using such models, and thus more tailored architectures to create ChEMBL wide bioactivity models.

## 3 Conclusions

We have created and benchmarked a standardized set based on high quality ChEMBL data (version 20). This dataset, together with the scripts used is available online, and can serve as a standardized dataset on which novel algorithms can be tested. Moreover, we have tested and compared a diverse group of established and more novel bioactivity modeling methods (descriptors, algorithms, and data formatting methods). To the best of our knowledge this is the first paper wherein in Deep Learning is coupled to proteochemometrics. Finally, we have explored the potential of DNNs by tuning their parameters and suggested ways for further improvement.

From our results we draw a number of conclusions. Most of the methods and algorithms can create models that are predictive (perform better than random). Training time versus accuracy is a less relevant issue as the best performing methods required less than 10 hours. Commonly used ‘random split’ partitioning might lead to an overly optimistic performance estimate. It is proposed to split training and tests sets based on time-based differences, providing a more challenging and more realistic performance estimate. It should also be noted that active and inactive compound-target combinations can impact the performance.

Focusing on machine learning methods, usage of DNN based models increases model prediction quality over existing methods such as RF, SVM, LR, and NB models. This is especially true when using multi task or PCM based DNNs and less so when using single task DNNs. As an added benefit, this gain in performance is not obtained at the expense of highly increased training times due to deployment of GPUs.

It was shown that the widest and deepest DNN architectures produced the best results in combination with the most descriptor features. There is certainly still room for improvement as hardware (memory) limitations or extreme training times were not reached. Moreover, model ensembles of the 63 individual models further enhanced the results yielding performance that was 27% better than the best performing model prior to tuning, indicating that indeed better results are possible.

Taken together, we anticipate that methods discussed in this paper can be applied on a routine basis and can be fine-tuned to the problem (e.g. target) of interest. Moreover, due to low training time and high performance we anticipate that DNNs will become a useful addition in the field of bioactivity modeling.

## 4 Methods

### 4.1 Dataset

Data was obtained from the ChEMBL database (version 20) [4], containing 13,488,513 data points. Activities were selected that met the following criteria: at least 30 compounds tested per protein and from at least 2 separate publications, assay confidence score of 9, ‘single protein’ target type, assigned pCHEMBL value, no flags on potential duplicate or data validity comment, and originating from scientific literature. Furthermore, data points with activity comments ‘not active’, ‘inactive’, ‘inconclusive’, and ‘undetermined’ were removed.

If multiple measurements for a ligand-receptor data point were present, the median value was chosen and duplicates were removed. This reduced the total number of data points to 314,767 (**Supplementary figure 2**), or approximately 2.5% of the total data in ChEMBL 20.

Typically, studies have used thresholds for activity between 5 and 6 [10, 11, 32, 33]. Data points here were assigned to the ‘active’ class if the pCHEMBL value was equal to or greater than 6.5 (corresponding to approximately 300 nM) and to the ‘inactive’ class if the pCHEMBL value was below 6.5. This threshold gave a good ratio between active and inactive compounds. Around 90% of the data points are active when a threshold of 10 μM is used, while a roughly equal partition (55%/45%) occurs at a threshold of 6.5 log units (**Supplementary figure 3**). Additionally, it represents an activity threshold that is more relevant for biological activity.

The final set consisted of 1,227 targets, 204,085 compounds, and 314,767 data points. Taken together this means the set 0.13 % complete (314,767 out of 250,412,295 data points measured). Moreover, on average a target has 256.5 (±427.4) tested compounds (median 98, with values between 1 and 4703).

ChEMBL L1 and L2 target class levels were investigated. For the L1 targets, most dominant are enzyme (144,934 data points) followed by membrane receptor (113,793 data points), and ion channel (16,023 data points). For the L2 targets G Protein-Coupled Receptors are most dominant (104,668 data points), followed by proteases (34,036 data points), and kinases (31,525 data points). See **Supplementary figure 7** for a graphical view. Finally, each compound has on average been tested on 1.5 (±1.3) targets (median 1, with values between 1 and 150). In total the set contained 70,167 Murcko scaffolds [39].

### 4.2 Compound Descriptors

Fingerprints used were RDKit Morgan fingerprints, with a radius of 3 bonds and a length of 256 bits. For every compound the following physicochemical descriptors were calculated: Partition Coefficient (AlogP) [40], Molecular Weight (MW), Hydrogen Bond Acceptors and Donors (HBA/HBD), Fractional Polar Surface Area (Fractional PSA) [41, 42], Rotatable Bonds (RTB). For descriptors used in the PP context please see **Supporting Information Methods**.

### 4.3 Protein Descriptors

For the PCM models, protein descriptors were calculated based on physicochemical properties of amino acids similar to previous work [43, 44]. However, lacking the ability to properly align all proteins, descriptors were made alignment independent which is different from our previous work. The sequence was split into 20 equal parts (where part length differed based on protein length). Per part, for every amino acid the following descriptors were calculated: Amount of stereo atoms, LogD [40], charge, hydrogen bond acceptors and hydrogen bond donors, rigidity, aromatic bonds, and molecular weight. Subsequently per part the mean value for each descriptor was calculated, and repeated for the whole protein, calculating the mean value for the full sequence length. Leading to an ensemble of 21 * 8 mean physicochemical property values (20 parts + global mean). Furthermore, sequence length was included as separate descriptor. It should be noted that this type of descriptor is a crude protein descriptor at best with significant room for improvement. However, the descriptor captures similarity and differences between the proteins and it is shown to improve model performance over models lacking this descriptor. Optimizing this descriptor is judged to be out of scope of the current work but planned for follow up.

### 4.4 Machine Learning – NB, RF, SVM, and LR models

Models were created using scikit-learn [45]. Naive Bayes models were trained using the same procedure as MultinomialNB [46]. A reference NB model with an activity threshold of 10 μM was included using PP and default setup.

RF were trained using the RandomForestClassifier. The following settings were used: 1000 trees, 30% of the features were randomly selected to choose the best splitting attribute from, with no limit on the maximum depth of the tree.

SVMs were trained using the SVC class, using the following settings: radial basis function kernel wherein gamma was set at 1 / number of descriptors. Further parameter cost was set at 1 and epsilon was set at 0.1.

For LR, the LR class of the linear_model package was used. The settings were mostly set to default, except for the solver, which was set to Stochastic Average Gradient descent with a maximum of 100 iterations.

### 4.5 Machine Learning – Neural Networks

In our experiments, we used a network with the following architecture: an input layer with, for example, 256 nodes representing 256 bit fingerprints, connected to 3 hidden layers of 4000, 2000, 1000 of rectified linear units (ReLU) and an output layer with as many nodes as the number of modeled targets (e.g. 1227 for the multi-task network). ReLU units are commonly used in DNNs since they are fast and unlike other functions do not suffer from a vanishing gradient. The output nodes used a linear activation function. Therefore, the original pChEMBL values were predicted and were subsequently converted to classes (pChEMBL ≥ 6.5 = active, pChEMBL < 6.5 = inactive). The target protein features and physicochemical features in the input layer were scaled to zero mean and unit variance. The output for a particular compound is often sparse, i.e. for most targets there will be no known activity. During training, only targets for which we have data were taken into account when computing the error function to update the weights. We chose to equally weight each target, for which we had data.

For training our networks we used stochastic gradient descent with Nesterov momentum which leads to faster convergence and reduced oscillations of weights [47]. Data was processed in batches of size of 128. After the neural network has seen all the training data, one epoch is completed and another epoch starts.

Moreover, after every epoch the momentum term was modified: the starting Nesterov momentum term was set to 0.8 and was set to 0.999 for the last epoch (scaled linearly). Likewise, during the first epoch the learning rate (the rate at which the parameters in the network are changed) was set to 0.005 and scaled to 0.0001 for the last epoch. These settings were decreased/increased on a schedule to allow for better convergence, increasing the momentum allows for escaping local minima while decreasing the learning rate decreases the chance of missing a (global) minimum.

To prevent overfitting of the networks, we used 25% dropout on the hidden layers together with early stopping [31]. The early stopping validates the loss on an evaluation set (20% of the training data) and stops training if the network does not improve on the evaluation set after 200 epochs (**Supplementary figure 7**). The maximum number of iterations was set to 2000 epochs. Dropout is a technique to prevent overfitting, by discarding, in each iteration of the training step, some randomly chosen nodes of the network.

To find the optimal network configuration we used grid search, limiting the number of epochs to 500 to speed up the training. In total, 63 individual models were trained to validate the influence of different descriptors, architecture and type of neural network (**Supplementary figure 6**). Due to problems with the stochastic gradient descent, for the PCM models with 4096 fingerprints plus physicochemical chemical properties, architectures 2, 4, 6 (**Supplementary figure 6**); a batch size of 256 instead of 128 was used. In all cases where physicochemical chemical properties were used, they were scaled to zero mean and unit variance.

In our experiments with neural networks we used nolearn/Lasagne and Theano packages [48-50] and GPU-accelerated hardware. The main script for training the networks is available in the supporting dataset deposit (‘FFNN.py’ and ‘instructions_DNN.pdf’).

### 4.6 Validation Metrics

We used the MCC and BEDROC as a primary validation metrics [26, 51, 52]. BEDROC (α=20), which corresponds to 80% of the score coming from the top 8% was used [26]. This was done to evaluate the performance in a prospective manner, where often the top % scoring hits is purchased.

Four separate MCCs and BEDROCs were calculated. One value for the pooled predictions (pooling true positives, false positives, true negatives, and false negatives) was calculated, and secondly an average MCC was calculated based on the MCC values per protein. Of these two values the mean is visualized in **figures 1** and **2**, with the unprocessed values given in **Supplementary tables 1** and **2**. This was done for both the random split set and temporal split set.

When no predictions were made for a given target-compound combination a random number was generated for this pair between 0 and 1. The reason for this is that we aimed to simulate a true use case and not cherry pick good or bad predictions. To be able to compare prediction quality across the different methods used random values were used, leading to MCC scores close to 0 for these cases. For values > 0.5 this score was deemed ‘active’ and anything below 0.5 was deemed ‘inactive’.

### 4.7 Validation Partitioning

Two different methods were applied to partition the data in training/validation sets. The first method that was used was a ‘random split’, herein 10% of the data was partitioned using semi-stratified random partitioning with a fixed random seed as implemented in PP and set apart for future reference. The remaining 90% was partitioned in the same way in a 70% training and 30% test set.

For the second method, a separate set was constructed wherein the year of the publication was the split criterion. All data points originating from publications that appeared prior to 2013 were used in the training set, while newer data points went into the validation set.

### 4.8 Hardware

Experiments were performed on a Linux server running CentOS 6.7. The server was equipped with two Xeon E5-2620v2 processors (hyperthreading disabled) and 128 GB RAM. GPUs installed are a single NVIDIA K40 and 2 NVIDIA K20s.

### 4.9 Software used

Python (version 2.7) was used with the following libraries: RDKit (version 2014.09.2) for the calculation of the fingerprints and descriptors [53], scikit-learn version 0.16 for the NB and RF [45]. For the neural networks we used Theano [54] and nolearn, together with Lasagne [49, 50]. For the supplementary information tables, Pipeline Pilot (version 9.2.0) [55], including the chemistry collection for calculation of descriptors, and R-statistics (R version 3.1.2) collection for machine learning [56] were used. Algorithms only reported in the supplementary information are mentioned in the supplementary information.

## 5 Abbreviations

AUC: Aurea under the curve
BEDROC: Boltzmann-enhanced discrimination of Receiver Operator Characteristic
DNN: Deep Neural Networks
IRV: Influence Relevance Voter
LR: Logistic Regression
MCC: Matthews Correlation Coefficient
NB: Naive Bayes
PCM: Proteochemometrics
PP: Pipeline Pilot
PS-IRV: Potency-Sensitive Influence Relevance Voter
PY: Python
QSAR: Quantitative Structure-Activity Relationship
RF: Random Forests
ROC: Receiver Operator Characteristic
SEM: Standard Error of the Mean
SVM: Support Vector Machines

## 6 Declarations

### 6.1 Acknowledgements

The authors would like to acknowledge NVIDIA and Mark Berger for generously contributing the GPUs. GvW would like to thank Sander Dieleman and Jan Schlüter for the fruitful discussion.

### 6.3 Author Contributions

EBL, GP, and GJPvW conceived the study. EBL, BB, NtD, and GJPvW performed the experimental work and analysis. All authors discussed the results, wrote, and proofread the paper.

### 6.4 Availability of data and materials

During review, the dataset supporting the conclusions of this article is available on SURFdrive; https://surfdrive.surf.nl/files/index.php/s/ktAy7GIdLXRvKHN It will be made available by deposition including DOI at 4TU.ResearchData.

### 6.6 Ethics approval and consent to participate

Not applicable

### 6.7 Consent for publication

Not applicable

